# Quantum Biology in Cellular Migration

**DOI:** 10.1101/2022.09.09.507322

**Authors:** Amy M. Vecheck, Cameron McNamee, Renee Reijo Pera, Robert J. Usselman

**Affiliations:** Chemistry Program, Department of Biomedical and Chemical Engineering and Sciences, Florida Institute of Technology, Melbourne, FL, 32904 United States; McLaughlin Research Institute, Great Falls, MT, 59405 United States; Department of Mathematics, California Institute of Technology, Pasadena, CA, 91125 United States

**Keywords:** quantum biology, radical pair mechanism, cellular magnetophoresis, image analysis

## Abstract

The impact of magnetic fields on cellular function is diverse but can be described at least in part by the Radical Pair Mechanism (RPM), where magnetic field intervention alters reactive oxygen species (ROS) populations and downstream cellular signaling. Here, cellular magnetophoresis within three-dimensional scaffolds was monitored in an applied oscillating 1.4 MHz radiofrequency (RF) magnetic field with an amplitude of 10 μT and a static 50 μT magnetic field. Given that cellular respiration or glycolysis can be increased based on the orientation of the RF magnetic field, this study focused on the parallel orientation to increase ATP synthesis. Results suggest that RF accelerated clustering and elongation after 1 day with increased levels of clustering and cellular linkage after 7 days. Electron microscopy provided additional topological information and verified the development of fibrous networks and extracellular matrix were visualized after 7 days in samples maintained in RF. Analysis of the distribution of cells within the scaffolds revealed that the clustering rate during the first day was increased nearly five times in the RF environment. This work demonstrates time-dependent cellular magnetophoresis that may be influenced by quantum biology (QB) processes and signaling that can further attenuate or enhance cellular bioenergetics and behavior.

## 1 Introduction

Some metabolic processes are suggested to utilize the radical pair mechanism (RPM) in the activation of molecular oxygen by reduced flavin enzymes to produce reactive oxygen species (ROS) (Usselman et al., 2014;Usselman et al., 2016). Spin-selective ROS signaling channels are responsible for altering biological systems at functional, cellular, or organism levels. This work investigates the time-dependence of radio frequency (RF) induced magnetophoresis of cells embedded within a natural polymer scaffold.

The RPM occurs between radical pair (RP) intermediates that can be initialized by either photo-activation or during light independent redox cycles (Steiner and Ulrich, 1989;Massey, 1994). RP spin dynamics can be impacted by weak static magnetic fields and oscillating resonance magnetic fields, affecting downstream biological chemical processes (Hogben et al., 2009). A key feature of the RPM is the coherent transition between singlet and triplet states and the ability to modulate singlet-triplet mixing by specific external magnetic fields. Within the paradigm of ROS and the RPM, singlet channels produce hydrogen peroxide, while triplet channels produce superoxide. Singlet-triplet modulation occurs when radiofrequency (RF) waves are applied to the RP that have sufficient coherent times, presumably > 1 μsec. Biological responses can be sensitive to RF resonance with appropriate static magnetic fields, where the point of intervention occurs from quantum coherent RPs initializing from normal biological processes.

In previous work, we measured hydrogen peroxide and superoxide levels with hyperfine resonance frequencies (7.0 MHz) and the resulting impact on cellular proliferation (Usselman et al., 2014). A subsequent study demonstrated an increase in mitochondrial respiration by parallel fields at Zeeman resonance (1.4 MHz), while increases in glycolysis have been observed in perpendicular fields (Usselman et al., 2016). Cellular bioenergetics was determined based on changes in oxygen consumption rate (mitochondrial respiration) and extracellular acidification rate (glycolysis). Both studies verified a relative partitioning of increased hydrogen peroxide and decreased superoxide levels, a hallmark triplet-born spin-correlated ROS radical pairs in the absence of visible light.

To further understand the role of RPM and ROS effects in cellular functions, we used low-field and low frequency magnetic resonance to monitor cellular cluster rates within a tissue engineering model. Cellular scaffolds under each condition provided information on scaffold degradation as a function of extracellular matrix growth (ECM) by use of Scanning Electron Microscopy (SEM). Based on the rate of gelatin degradation, acellular scaffolds were estimated to lose shape during a short-term study of 7 days; without the addition of ECM, there is not enough support material to maintain the construct. The addition of RF was hypothesized to enable cells to create more ECM, which was visualized as a mesh-like material over and around scaffold pores.

## 2 Materials and Methods

### 2.1 Cell Culture

NIH3T3, mouse fibroblasts, were seeded at a density of 6,600/cm^2^ per T-75 flask containing 10 mL of media. All flasks were maintained in an incubation unit at 37 °C and 5% CO_2_. Cell culture media was changed every 2 days. When each flask reached between 85% and 90% confluency, media was aspirated into a collection beaker using a serological pipette. Phosphate Buffer Solution (PBS) was added to the flask to wash the lifted or dead cells and immediately aspirated into the collection beaker. Cell were trypsinized for 5 minutes and neutralized using an equal volume of media. Using a serological pipette, the contents of the flask were moved to a 15 mL conical tube and centrifuged at 1300 rpm for 5 minutes. Without disturbing the resulting pellet, the supernatant was aspirated into a collection beaker. Using a hemocytometer, cells were counted and split between new sterile flasks, each containing 10 mL of media.

### 2.2 Bioink Synthesis

Alginate-gelatin hydrogels were synthesized in a 5% (weight/volume) composition at a rate of 2:3 respectively. In a beaker with 10 mL deionized water and 10 mL PBS, 0.2 g sodium alginate was added gradually to the solution, which was spun at 350 rpm for 10 minutes to prevent clumping. The solution was heated to 40 °C to allow the added 0.3 g of gelatin to mix properly. In a 3 mL syringe, 1 mL of the alginate-gelatin mixture was aspirated, and the syringe was attached to a stopcock. In a second 3 mL syringe, 200 μL of cell culture media was aspirated, and the syringe was attached to the opposite end of the stopcock. For cellular constructs, a cell density of 2 × 10^6^ cells/mL was mixed into the media prior to aspiration. Plunges were pressed in an alternating pattern to homogenously mix the solutions. The entirety of the solution was maintained in one syringe until the extrusion process.

### 2.3 Bioprinting-Based Construct Formation

Constructs were formed using an extrusion-based methodology to distribute bioink in a layer-by-layer fabrication method. This methodology did not alter the mechanical properties of the construct as previously demonstrated (Kharel et al., 2019). However, the porosity of the sample would not be homogenous, as the resolution of the print is not exact during the hand-extrusion process.

Using the 3 mL syringe filled during the bioink synthesis stage, 1 mL of bioink was extruded in a 15 mm x 15 mm x 4 mm polylactic acid mold, which was printed using a Creality Ender 3 printer. Extrusion was completed over 10 seconds to minimize shear stress on the cells. The top of the construct was lightly coated with 0.5 M CaCl2 to prevent the alginate from beading during crosslinking. Full crosslinking was conducted by submerging the print in 0.5 M CaCl2 for 10 minutes. All constructs were maintained in either a static 50 μT magnetic field control or a static 50 μT and 1.4 MHz magnetic field until tested.

### 2.4 Magnetic Field Instrumentation

A tri-axial Helmholtz coil system with a 6-channel DC power supply was used to control the 50 μT static magnetic field direction in each of two incubators. A free-scale chip provided data for a feedback sensor that allowed for real-time control of magnetic fields within the incubators to cancel out stray fields. The static magnetic field noise floor is approximately 20 nT. A secondary Helmholtz coil was used to create an oscillating magnetic field at 1.4 MHz. While the secondary coils existed in both incubators, they were only powered for one of the incubators, which contained RF samples. The orientation of the RF was parallel to the static magnetic field and in-plane of the cell wells. This was set to roughly 10 μT_RMS_ amplitude, parallel to the static magnetic field.

### 2.5 DAPI-Phalloidin

Cellular constructs were maintained in either a 50 μT static field or 50 μT static field with 1.4 MHz RF for 0, 1, or 7 days. At such time points, constructs were washed with PBS and submerged in 3.7% formaldehyde for 15 minutes to fix the cells. The constructs were again washed with PBS and submerged in 0.1% Triton-X, a permeabilization buffer, for another 15 minutes. Samples were again washed with PBS and submerged in phalloidin for 1 hour prior to imaging. Samples were tested using confocal microscopy.

### 2.6 Scanning Electron Microscopy

Acellular and cellular constructs were tested on day 0, 1, and 7 to demonstrate the porosity and support materials of the samples as a function of time. All samples were dried in ethanol solutions, which increased by 10% every 10 minutes. Prior to this step, cellular samples were fixed with 3.7 % formaldehyde for 1 hour. Critical point drying was conducted to replace the ethanol with carbon dioxide. Samples were gold sputter coated for 60 seconds. A second round of sputter coating occurred after rotating samples approximately 90°; this was done to ensure full coverage of the sample. Acceleration voltage for the SEM was set between 5 kV and 10 kV.

### 2.7 Image Processing

Image processing was done to identify the locations of cells in images to determine clustering. Images were processed using MATLAB. First, the image was converted to a binary image with a threshold value of 0.1. The images were then morphologically opened and closed using reconstruction to produce a noiseless image. The images were then segmented using watershed segmentation. The centroids of the newly segmented regions were identified and labeled. These labels were assumed to be the locations of the cells in the images. The coordinates of each centroid were stored and used for data analysis (Supplementary Methods).

### 2.8 Data Analysis

Using centroids generated through image processing, change in cell density over time was determined. First, the distribution about the geometric center was calculated. The collected data was then normalized relative to the number of cells in the tissue. Using a cumulative distribution histogram, a degree four polynomial function was fitted to the curve. This polynomial was used to determine the half-radius (R50) of the cells in the image. This was done by solving for the polynomial when y=0.5. This value was taken as R50. Cell density was then determined by dividing half the number of cells identified by the area encompassed within the determined R50 for that image (Table 1). Density was calculated in cells per micrometer squared. The cell densities were then compared for days 0, 1, and 7 to create a change in density over change in time (Supplementary Methods).

**Table 1:**
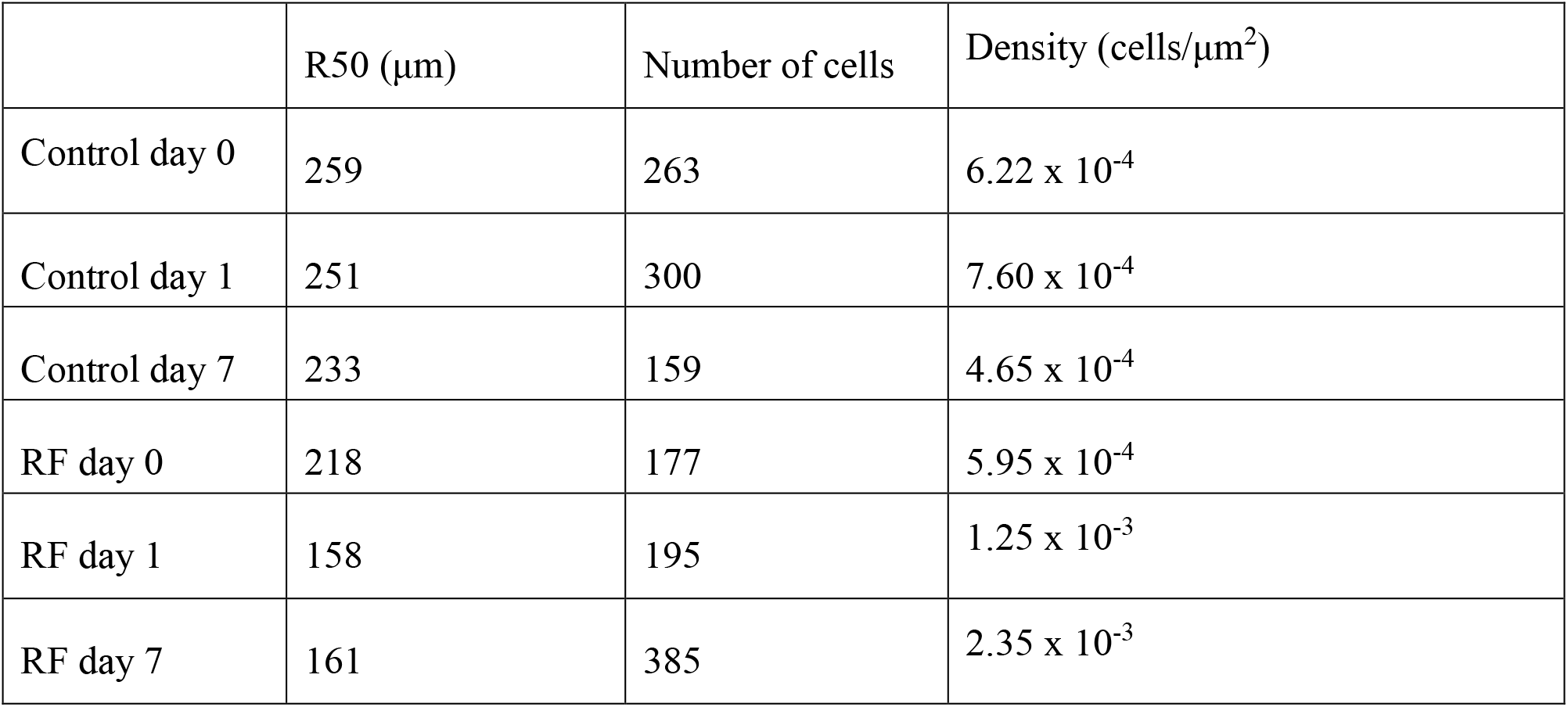
Comparing Cell Densities.

## 3 Results

### 3.1 Orientation of the Cytoskeleton

DAPI/phalloidin staining was used to label the cells’ nuclei and cytoskeleton, respectively, as depicted in Figure 1 and Figure 2. Day 0 samples showed a homogenous distribution of the cells throughout the construct, which was expected based on the results of the live/dead assay. Day 1 samples maintained in a static 50 μT magnetic field yielded slightly more clustered results. Day 7 clustering results are presumably within the variation of cellular seeding for different sample days. These were smaller clusters than seen in the day 1 samples maintained in RF.

**Figure 1:**
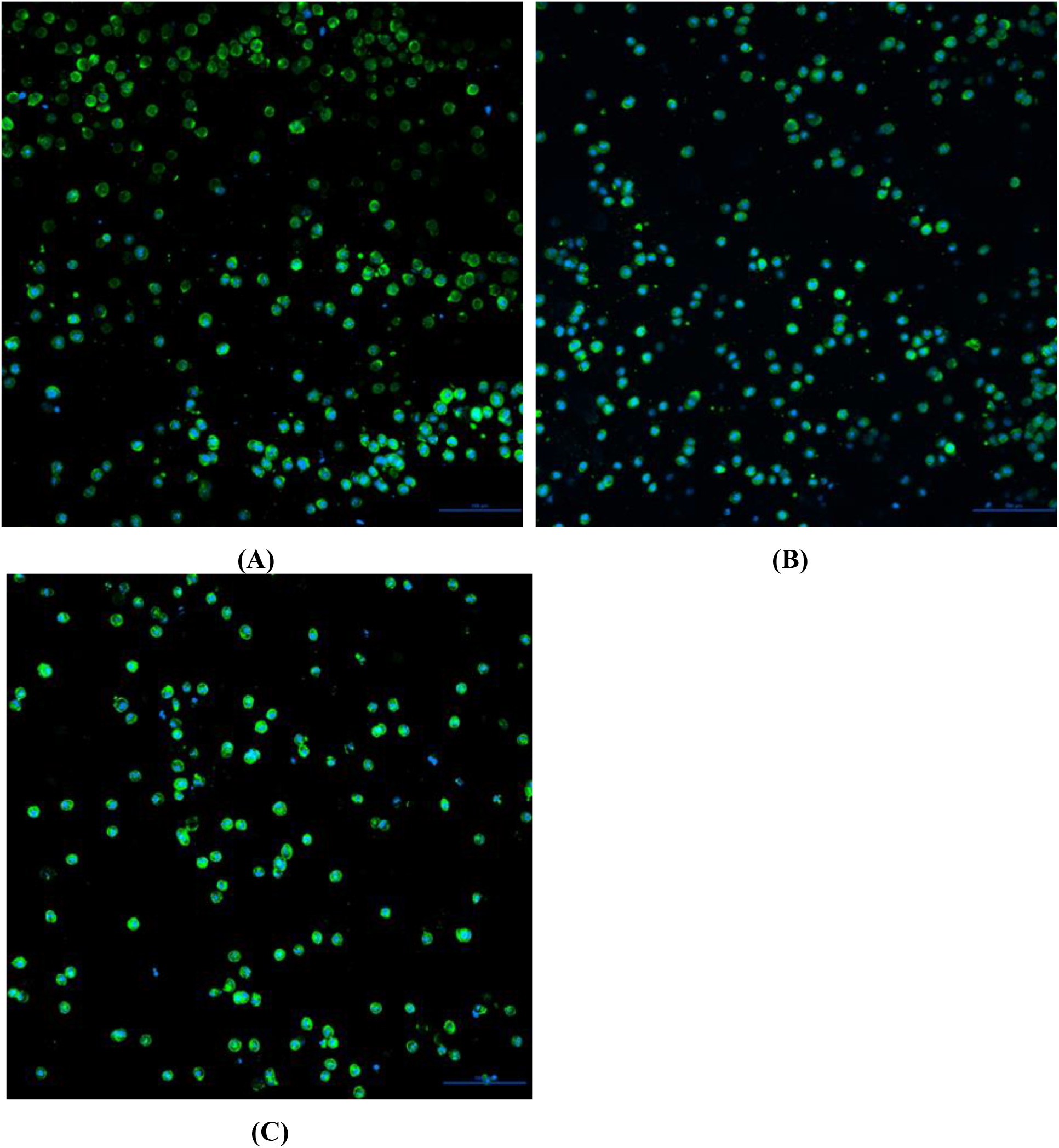
Nuclei and cytoskeleton staining demonstrating the distribution of cells **(A)** initially after construct synthesis, **(B)** after 1 day at 50 μT, and **(C)** after 7 days at 50 μT

**Figure 2:**
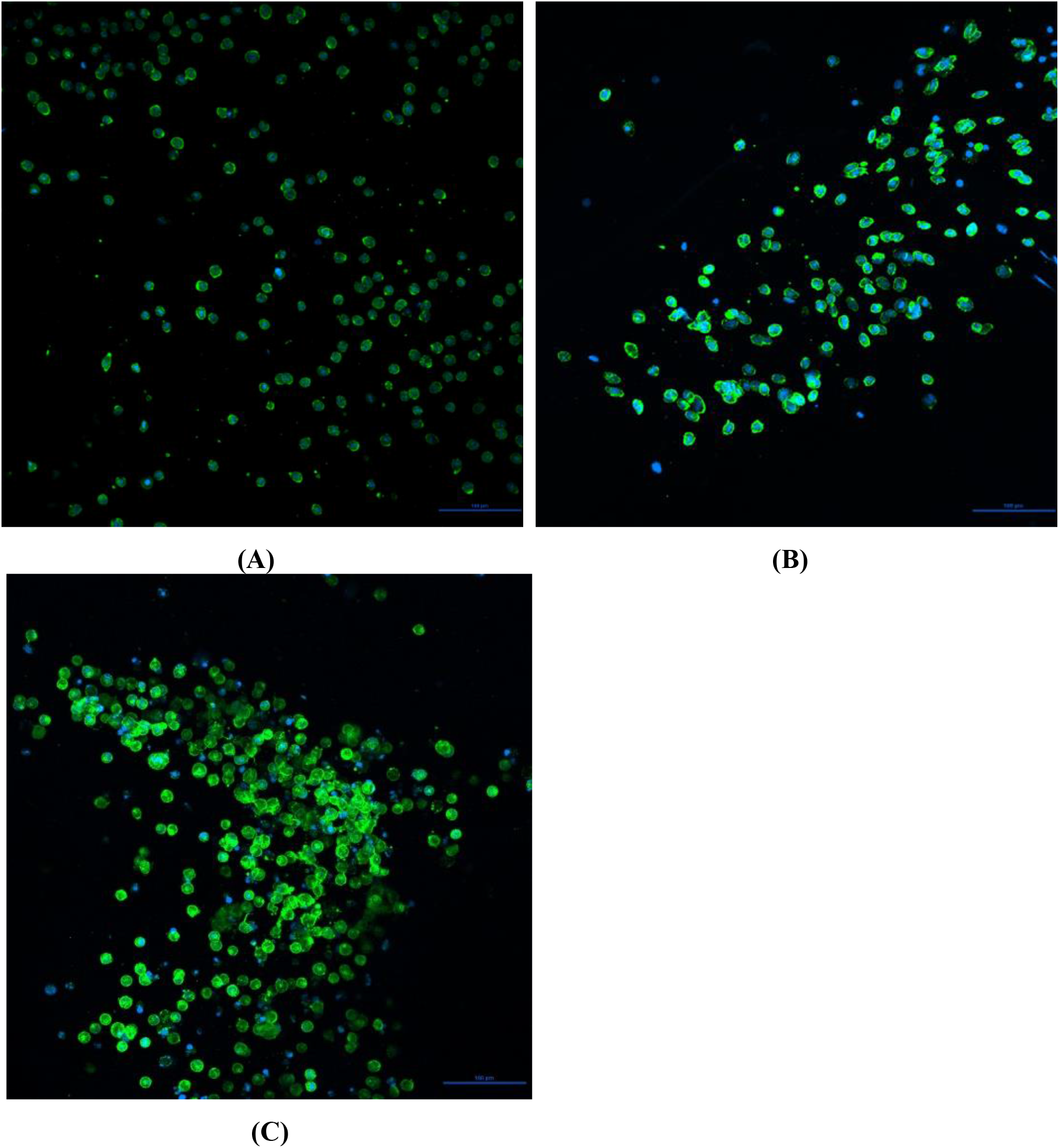
Nuclei and cytoskeleton staining demonstrating the distribution of cells **(A)** initially after construct synthesis, **(B)** after 1 day at 50 μT + RF and **(C)** after 7 days at 50 μT + RF

**Figure 3:**
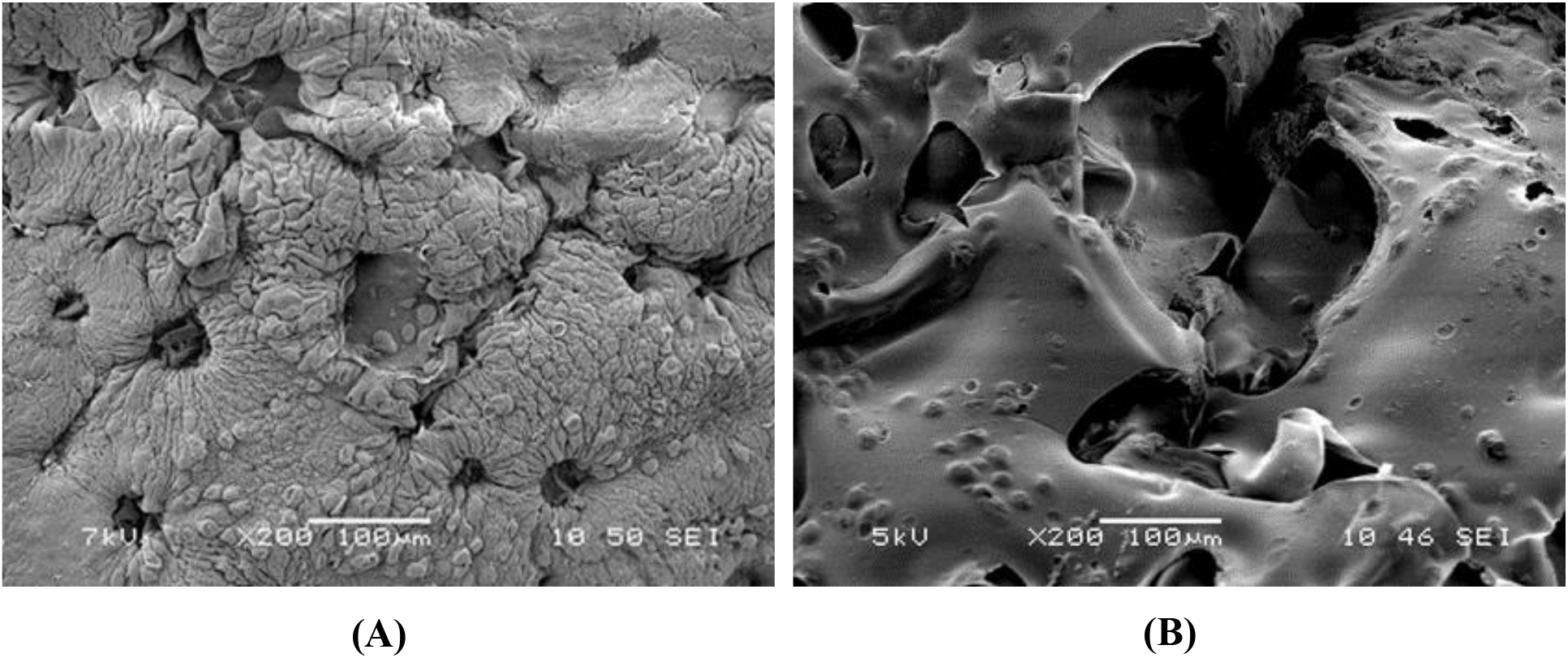
Sample SEM images taken at 200X of **(A)** cellular constructs at day 1, and **(B)** cellular constructs at day 7. All constructs shown were maintain in a static 50 μT magnetic field

Migration was determined based on characteristic elongation of the cells. Cells did not show elongation in the same direction as is expected in standard cultures. Due to the internal structure of 3D constructs, this type of standardized elongation is not possible. Rather, the cells will stretch around pores enabling nutrient flow and easy communication with other cells. The extent of the cellular signaling was not able to be determined within the scope of this study. The clustering for RF maintained samples continued occurring over the course of the seven days. Multiple sections of each sampled construct showed that the clustering occurred throughout, but elongation ceased once the cytoskeleton of multiple cells began to overlap. The cell linkage seen for the day 7 samples with the addition of RF show similar results to previous testing of day 10 samples with fibrinogen included in the bioink. For this reason, it can be concluded that the addition of RF increases cellular mobility and cell-cell signaling. Increased cellular migration rates could be due to an oxidative signaling adaptive response (Sies et al., 2022), facilitated by ROS-mediated Zeeman resonance, affecting mitochondrial bioenergetics and cytoskeletal movement.

### 3.2 Surface Topography and ECM Formation

The majority of the open cell pores appear to exhibit tearing of the biomaterial. Alginate is commonly used to synthesize microbeads due to its ability to encapsulate its surroundings. The drying process can explain the rupture based on this as the removal of water using ethanol could have caused any surface beading of the alginate to break apart as the fluids exchanged. Cellular constructs showed less rupture of the surface; however, both open cell and closed cell pores were less homogenously distributed. This may have been caused by the distribution of cells along the surface of the material.

SEM of constructs maintained in a 50 μT static field demonstrated the basic thermos-reversible properties of gelatin as well as the structural support that cells provide; these samples can be seen in Figure 4. Acellular constructs after 1 day in incubation demonstrated a significant increase in porosity. A large portion of gelatin dissolved into the PBS for all samples as previously hypothesized. No acellular constructs were viable by day 7 as they did not maintain sufficient structural support to withstand the drying process. Cellular constructs on day 1 showed an increase in the number of cells on the scaffold surface. While scaffold degradation did occur and open cell pores were created more homogenously, the degradation was not as drastic as the acellular samples. This is because cells have attached to the binding sites of the gelatin and cross-linked it in a way that prevented it from easily dissolving into the PBS. Day 7 samples yielded similar results, but with a more significant surface area due to open cell pores.

**Figure 4:**
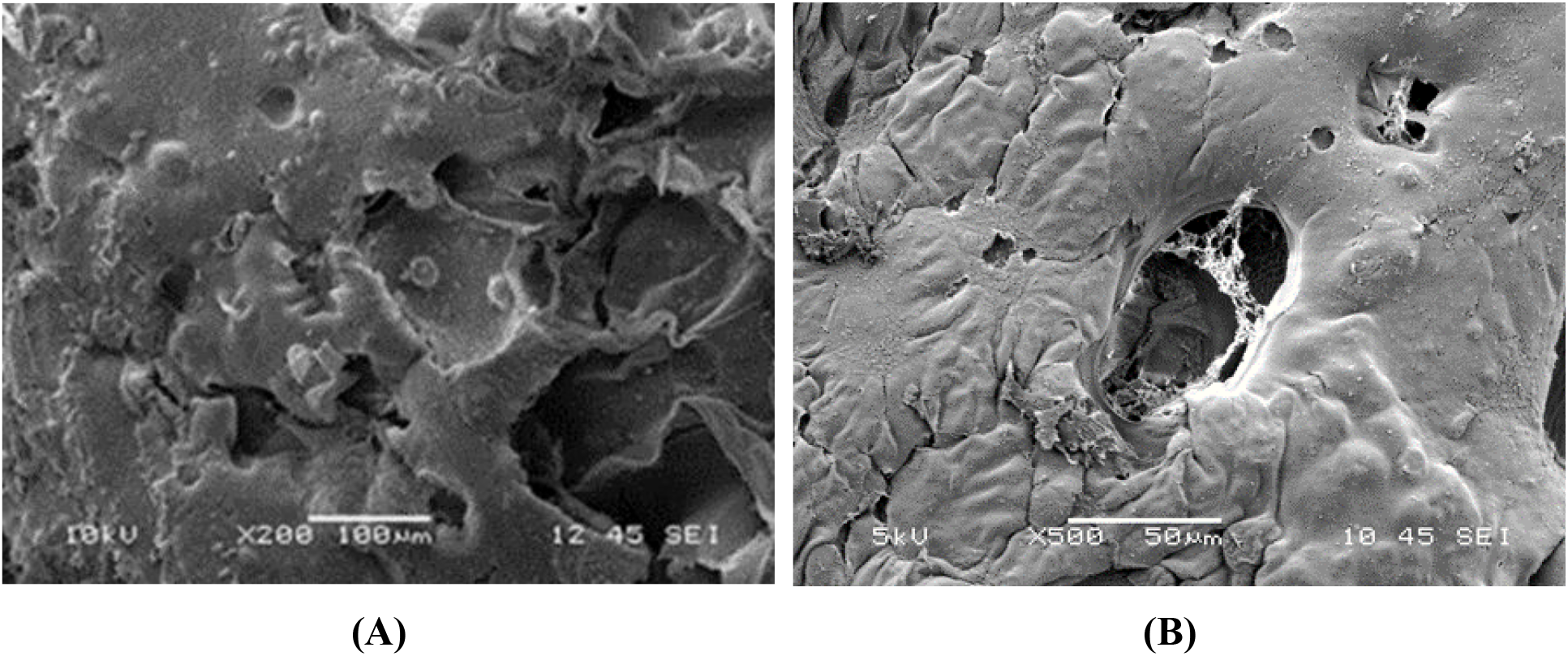
Sample SEM images taken between 200X and 500X of **(A)** cellular constructs at day 1, and **(B)** cellular constructs at day 7. All constructs shown were maintained in a static 50 μT magnetic field with 1.4 MHz

SEM of constructs maintained in a 50 μT static field with the addition of 1.4 MHz demonstrated an increase in porosity for both acellular and cellular constructs as seen in Figure 4. Acellular constructs in RF dissolved significantly by day 1, which created an unstable surface for the sputter coating. While the construct maintained its general shape, the irregular topography enabled a significant amount of sun-spotting for the images and caused damage to all samples. Day 7 acellular constructs were not viable for the drying process as they had dissolved into globules. Day 1 cellular constructs showed increased porosity, specifically for closed cell pores. Fixed cells appeared to be larger in size, which was confirmed previously with live/dead assay and DAPI/phalloidin staining. Day 7 cellular samples showed cells located around the pores with mesh-like networks commonly overlaying pores less than 40 μm. These mesh-like networks are indicative of fibers being deposited by the cells and can be considered part of the ECM. In longer duration studies, the addition of the ECM is expected to begin to cause gelatin to dissipate more readily in cellular constructs.

### 3.3 Change in Cell Density

Clustering rate was determined for both environments at initial seeding, after 1 day of treatment, and after 7 days of treatment (Figure 5). A novel method was developed that determines changes in cell density over time. This method allows for the global distribution of the cells to be captured and uses this distribution to describe clustering. The comparison of cell density over time determines the clustering rate. This gives a metric for total migration of cells within a sample. It is this method the allows one to transition from visual inspection of cell images to comparing mathematical measurements that accurately describe the migration of these cells. This provides us with a deeper understanding of the cell migration time-dependence and gives us a route for direct numerical comparison between samples. Without this novel method, any data depicted about cell clustering would be purely qualitative. The novel image analysis provides a quantitative method to measure differences in cell migration rates.

**Figure 5:**
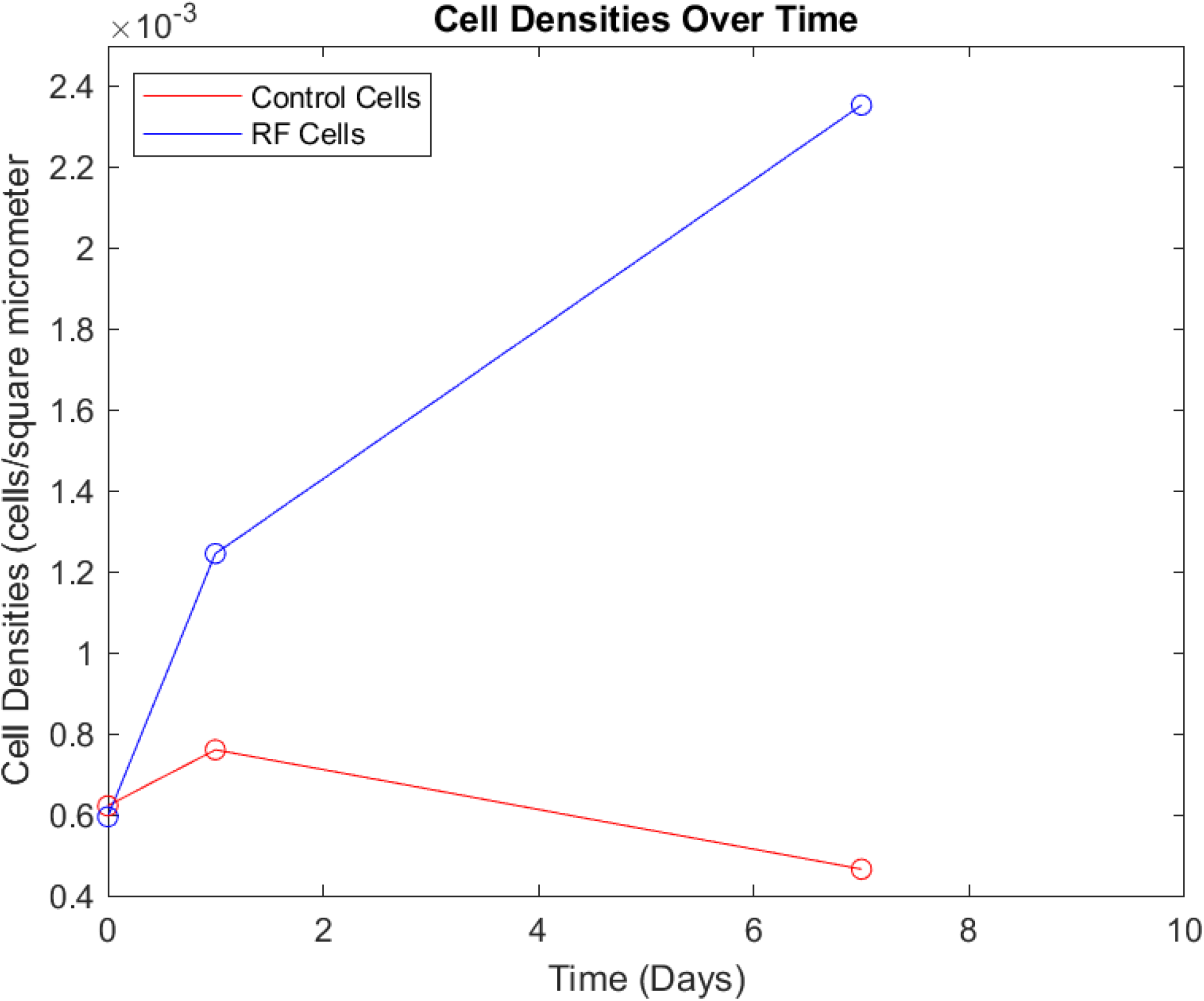
Cell densities calculated over time. Both start at nearly the same cell density, which is expected because we predict a homogenous initial seeding.

The cells maintained in RF had a rate of 6.55 × 10^−4^ cells·μm^−2^, as compared to the control cells with a decreased rate of 1.38 × 10^−4^ cells·μm^−2^ per day (Table 1) from initial seeding to day 1. This result suggests that the RF treatment led to a clustering rate 4.7 times that of the standard tissue during the first day of treatment. From day 1 to day 7, the RF tissue slowed clustering with a rate of 1.83 × 10^−4^ cells·μm^−2^ per day. The control cells appeared less clustered during this time with a rate of −4.92 × 10^−5^ cell·μm^−2^ per day (Figure 5). This suggests that RF continues to increase clustering from the first day to day seven but does so at a slower rate as compared to the first day.

## 4 Discussion

This study assessed the effects of cellular migration by modulating the RPM using Zeeman resonance. Cells were added to a 2:3 alginate-gelatin hydrogel to for a 5% w/v bioink. All samples were extruded using a bioprinting-based methodology and casted in a PLA mold. After crosslinking with 0.5 M CaCl2, constructs were maintained in either a 50 μT static magnetic field or a 50 μT static field with RF. Cellular viability assays verified that there were no cytotoxic effects of the biomaterials when maintained in a 1.4 MHz magnetic field. This assay gave an indication of cellular migration due to clustering in day 1 samples maintained in RF. Nuclei and cytoskeleton staining provided data regarding clustering patterns such that RF samples showed clustering and elongation after 1 day and showed higher levels of clustering and cellular linkage after 7 days.

SEM provided indications of scaffold degradation including that acellular sample degradation occurred faster in RF environments. Fibrous networks were noted in pores for cellular samples maintained with RF for 7 days. It is believed that the increase in basal respiration, due to utilization of the RPM, increased ATP production, thus increasing migration and cellular signaling. From our data analysis, we were able to determine the clustering rate over time for the RF treated tissues and the control tissues. Through a direct comparison of these rates, we calculated that the RF treated tissue clustered at a rate 5.8 times faster than the control tissue during the first day of treatment and continued to increase clustering over seven days of treatment. This suggests that RF dramatically increases the migration rate of cells, allowing them to not only cluster much more, but do so faster, reaffirming our hypothesis that parallel RF magnetic fields increase cell performance.

Concepts of quantum biology (QB) began approximately 8 decades ago (McFadden and Al-Khalili, 2018) and aim to observe nontrivial, quantum signatures written into biochemical mechanisms that impact cellular function. One such mechanism is the RPM (Hogben et al., 2009) (Zadeh-Haghighi and Simon, 2022). Since the 1970s, the RPM has been used to describe the magnetic field effects on bird navigation (Hore and Mouritsen, 2016). The physiological magnetic compass of birds allows them to make directional and positional decisions in flight. Klaus Schulten first hypothesized the radical pair mechanism as the basis for such navigation (Schulten et al., 1978;Schulten and Wolynes, 1978). Outside of magnetic fields, migratory birds have also shown a photosensitive dependance for navigation (Mouritsen, 2018). This has been attributed to cryptochromes, a photo-magnetoreceptor molecule found in migratory birds and plants (Ahmad and Cashmore, 1993). Enhanced cryptochrome response varies as it affects the circadian clock in animals but additionally increases the development and growth responses of plants (Arthaut et al., 2017;Hammad et al., 2020). The RPM quantum effects in broader areas in ROS redox biology are emerging (Usselman et al., 2016;Pooam et al., 2020). Discovering how we can influence cellular systems using QB principles is at the core of this study.

Further work to support our hypothesis would involve direct imaging of ROS to gain more insight into the mechanism involved. This would give us a deeper understanding of RPM and the impact on biological function. There are many options for this imaging technology, many of which are effective for even extremely low ROS concentrations (Shen et al., 2020). In addition, observing ROS species-specific measurements would lend us quantitative information as to the population changes due to altered spin dynamics. Fluorescent and electrochemical options to date are extremely effective for this kind of approach (Bai et al., 2019;Duanghathaipornsuk et al., 2021).

Physiological applications for ROS and QB exist in not only tissue engineering but also neurodegeneration and aging models. Because of parallel RF oscillating magnetic field’s ability to modulate the production of ROS, there is an avenue for treating certain diseases such as Alzheimer’s Disease (AD). Amyloid β plaques are a well-known characteristic of AD, but high levels of ROS accompany and are thought to precede these plaques (Huang, Zhang and Chen, 2016; Angelova and Abramov, 2018). Superoxide is the primary source of oxidative stress in the brain and leads to downstream damage in nervous system tissues (Huang et al., 2016;Angelova and Abramov, 2018). A possible way to alleviate oxidative stress in the brains of AD patients lies in the methods given by this paper. We have shown that RF has notable and dramatic effects on cell performance and predict that we can use these effects to treat diseased neurons. The applications are waiting to be tapped into. Antioxidant therapies have shown lots of promise for the slowing of AD. We could analyze the change in superoxide production using specific ROS sensors, allowing us to provide accurate measurements for efficacy. This treatment could provide more specific and timely results with limited adverse side effects, as compared to traditional methods of treatment.

More generally, evidence is accumulating that supports the RPM involvement in altering ROS levels and signaling channels in redox cellular processes (Usselman et al., 2016;Pooam et al., 2020;Bradlaugh et al., 2021;Ikeya and Woodward, 2021;Deviers et al., 2022). A novel area in ROS QB is emerging that connects the RPM at the quantum level to the classical spatial-temporal domains in living systems, across electron spin dynamics, flavoenzymes, mitochondria bioenergetics, cytoskeleton, and ultimately cellular function (Figure 6). We propose a novel framework that introduces a type of Quantum Biology Clock (QBC) that impacts select spatial-temporal domains with normal cellular function in synchronous modes. Magnetic resonances alter the singlet-triplet mixing at the point of ROS generation, thus changing the timing of the QBC in an asynchronous fashion. The cells respond to the QBC by adapting to the activated oxidative signaling channels that percolate through the spatial-temporal domains. This work represents a manifestation of ROS QB and supports evidence of a novel QBC operating as a control system that originates at the quantum level.

**Figure 6:**
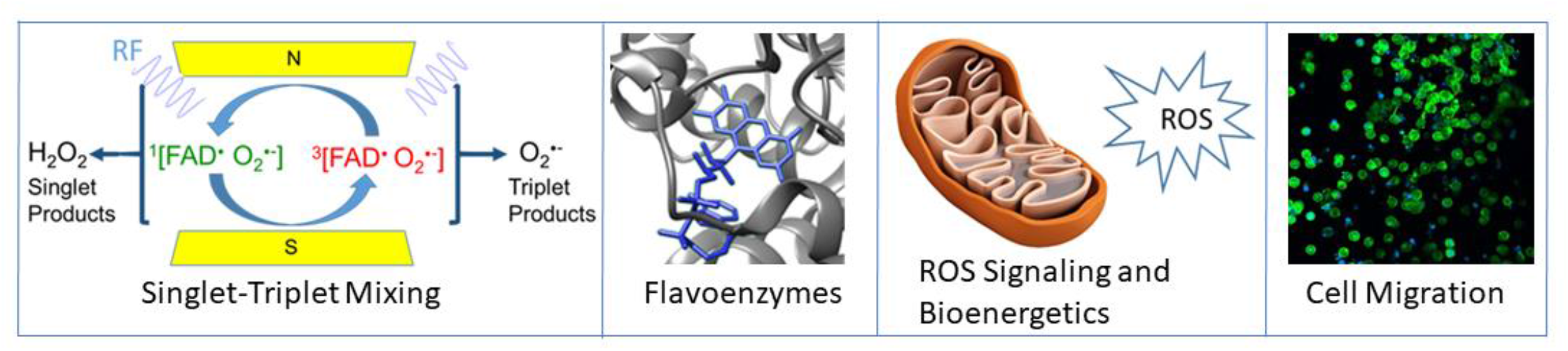
Spatial temporal domains that link the timing of a QBC in living systems.

## 2 Author Contributions

R.J.U. conceived of the presented idea. A.M.V. wrote the manuscript with support from R.J.U., generated the cell tissues, and provided the cell images. C.M. wrote the manuscript with support from R.R.P. and R.J.U., provided data for Table 1, and conducted all data analysis and computation. All authors contributed to the final manuscript.

## 4 Acknowledgments

R.J.U. would like to thank members of his research group for helpful comments on the manuscript.

## 1 Data Availability Statement

The datasets for this study can be found in github repository at https://github.com/Cameron-McNamee/Cluster-Analyzer

