## Supplemental Materials for "Quantum Biology in Cellular Migration"

### *Supplementary Material*

#### 1 Supplementary Methods

##### 1.1 Image Processing

The raw images were first edited to be 750-by-750 pixels (Fig. 1A). This was done to ensure consistency across all images and to improve accuracy. These dimensions were found to be optimal for providing the highest image quality while allowing the algorithm to run with the highest speed. Higher pixel density images were attempted, and they revealed levels of noise that could not be removed without also removing substantial portions of signal. Lower pixel density images lead to inaccuracies with the image processing, causing the algorithm to misidentify cells because of the high loss in the noise removal process.

Images were then converted to grayscale (Fig. 1B). This was done as an intermediate step towards binarizing the images. The grayscale images were then binarized using a luminance threshold of 0.1 (Fig. 1C). Otsu's method was attempted, along with an adaptive threshold. Otsu's method was not detailed enough and lead high amounts of loss for fainter cells. This is likely due to the low difference between the foreground and background in certain spots. An adaptive threshold picked up extreme amounts of noise, leading to entire sections of images coming up as signal when there were no cells present.

The binary image was morphologically eroded by a disk of radius five. The eroded image was then used as a marker and the original binary image as a mask for morphological reconstruction. This process effectively completed morphological opening. Opening was tried directly, but it removed too much signal and yielded an image that was far too different from the original. Reconstruction has the added advantage of producing an image that is more like the original, leading to higher accuracy and lower loss.

The opened image was then morphologically dilated using a disk of radius five. The compliment of the dilated image was used as a marker and the compliment of the opened image as a mask for morphological reconstruction. The compliment of the image produced was then taken. Similarly, this process was effectively morphological closing. For the same reasons outlined above, closing was not used directly (Fig. 1D).

After noise removal, the image was segmented using watershed segmentation (Fig. 1E). The reason for segmenting was to produce a more accurate dataset that identified each cell. This process was attempted without segmentation, and it was unable to identify closely linked cells that were obviously not individual cells. With the addition of segmentation, the datasets were much more representative of the raw images. The watershed segmentation was done by first converting the modified image to a distance transform. The compliment of this transform was used to create the watershed. The pixels that were background in the binary image were set to zero in the watershed. This produced a labeled image that could then be used for identifying the individual cells.

The centroids of each label were found and stored as the locations of the cells (Fig. 1F). This completes the image processing.

**1.2 Data Analysis**

Once the cells were identified, the distribution was able to be determined. This was done by first identifying the geometric center of the cells (Fig. 2A),

$$x_{mean} = \frac{\sum_{c \in C} c_x}{|C|}$$

$$y_{mean} = \frac{\sum_{c \in C} c_y}{|C|}$$

where *C* is the set of all cells, $c_x$ and $c_y$ are the x- and y-coordinates of a given cell *c*. With this as the reference point, the distance of each cell to this center was then calculated.

$$d_c = \sqrt{(x_{mean} - c_x)^2 + (y_{mean} - c_y)^2}; \ \forall c \in C$$

With these distances, a normalized histogram was produced. This showed the relative number of cells within a given radius (Fig. 2B). Using this graph, a degree four polynomial was fitted to the curve (Fig. 2C). This x-values of the polynomial when equal to 0.5 were then found.

$$p_4 x^4 + p_3 x^3 + p_2 x^2 + p_1 x + p_0 = 0.5$$

Imaginary and non-positive roots were excluded. This was called the half-radius or R50. This radius was the minimum radius required to produce a circle, centered about the geometric center, to include half the total number of cells in the image (Fig. 2A). The reasons for calculating the radius containing half the total number of cells, rather than some other amount, were: 1) having a radius dependent on a smaller proportion of the cells did not accurately depict the entire population of the tissue. This was because a smaller radius would be highly dependent on the cells surrounding the immediate area about the geometric center. This gives a poor metric of the entire picture. 2) A radius including a higher proportion of the cells would be highly dependent on outliers. This would cause images with cells around the border to have disproportionately high R50's, which is not an accurate measure of clustering.

With the R50 calculated, cell density was determined.

$$\rho = \frac{|C|}{2\pi (r_{50})^2}$$

This gives a density in cells per mircometer$^2$. The reason for having 2 in the denominator is to divide the number of cells in half. This way, we consider half the number of cells over the area needed to cover the same number of cells. This does not make a difference when comparing the rates between one another, but nevertheless gives more sensible values that correspond with what is being done in theory. With this calculated density, we then compared the different images over time to determine the clustering rate.

$$\frac{\rho_a - \rho_b}{a - b} = \frac{\Delta\rho}{\Delta t}$$

With a direct comparison of these calculated rates, we were able to show the increased clustering rates that supported our hypothesis.

#### 2 Supplementary Figures

##### 2.1 Figure 1

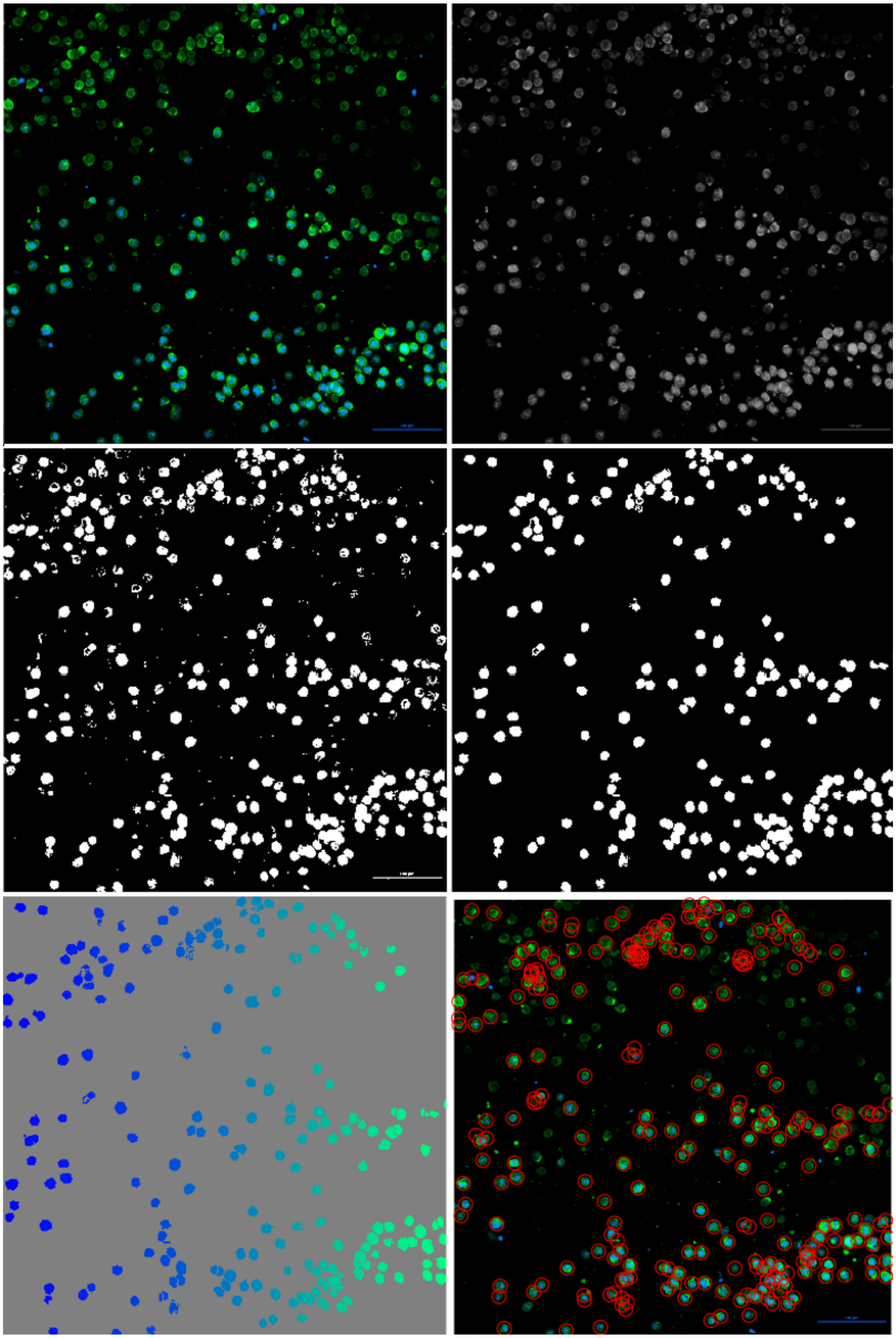

**Supplementary Figure 1.** (From left to right, top to bottom) The image used here is the control tissue upon initial seeding. **(A)** The raw image that has been converted to 750-by-750 pixels **(B)** The image once it has been converted to grayscale **(C)** The grayscale image after being converted to a binary image **(D)** The binary image after undergoing the noise removal process **(E)** The labeled map produced after segmenting the noise-removed binary image using watershed segmentation **(F)** The cells identified by the centroids of the labels superimposed on the original image. Here, one can see the robustness of the algorithm. Picking up faint cells that may be identified with the naked eye is not plausible without also picking up on high amounts of background noise.

**2.2 Figure 2**

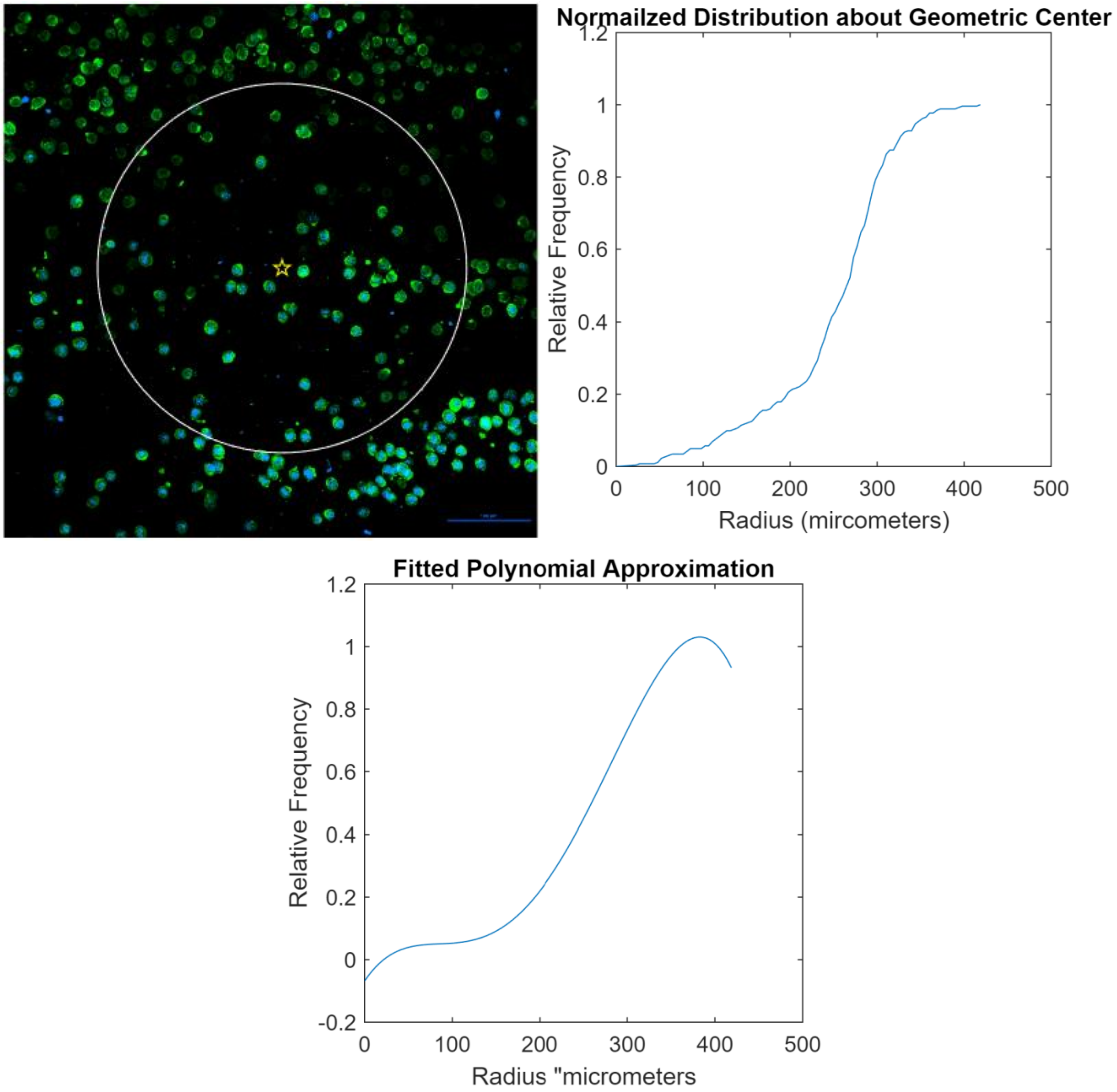

**Supplementary Figure 2.** (From left to right, top to bottom) The image used here is the same as above, control tissue upon initial seeding. **(A)** The geometric center drawn as the gold pentagram near the center and the circle described by the calculated R50 for this image. **(B)** The graph of the distribution about the geometric center. For a given radius (x), the relative frequency is the corresponding value (y). Here, relative means the number of cells within a radius divided by the total number of cells. This means when y=0.5, the radius encompasses half the cells. **(C)** A degree four polynomial fitted to the curve described in **(B)**.
